# Rapid generation and screening of transgenic black soldier fly (*Hermetia illucens*)

**DOI:** 10.1101/2024.02.21.581498

**Authors:** Chandran Pfitzner, Kate Tepper, Sheemal Kumar, Carly Retief, Justin M McNab, Robert A Harrell, Maciej Maselko

## Abstract

**Background:** The black soldier fly (BSF), *Hermetia illucens* is a widely used, and mass-produced insect that fulfils an important role in both the management of organic waste and as a component of animal feed formulations. They also have significant potential as a platform for converting organic waste into high-value proteins, and lipids for the production of biofuels. Applying synthetic biology to BSF provides even more potential for improvement through the generation of transgenic BSF to enhance animal feed, produce and fine tune high-value industrial biomolecules, and to expand their waste conversion capabilities.

**Results:** To enable the rapid generation and screening of transgenic BSF, we utilised microinjections of piggyBac mRNA with donor plasmids. We have found preliminary screening of G0 BSF to identify mosaics for outcrossing can be completed less than 2 weeks after microinjection. Stable transgenic lines were reliably generated with effective transformation rates of 30-33%, and transmission of the transgene could be confirmed 3 days after outcrossing the G0 adults. We also present a protocol for identifying the location of integrated transgenes.

**Conclusions:** The methods presented here expedite the screening process for BSF transgenesis and further expand the toolkit for BSF synthetic biology.

## Background

The black soldier fly (BSF), *Hermetia illucens*, has increasingly been recognised for its impressive dietary adaptability. This has quickly positioned BSF as a crucial organism for addressing the challenge of handling the 880 million tons of organic waste generated worldwide each year.^1^ By utilising this organic waste as their primary food source, BSF larvae become a valuable product. They are currently being used as a sustainable feed ingredient for animals such as fish, swine, and poultry,^2, 3^ and the processed organic waste, frass, can be re-used as a fertiliser to promote crop growth.^4^

The pressing concern for solutions to reduce the effects of climate change has prompted the need to transition from fossil fuels to cleaner, renewable energy sources. BSF offers another solution here as the lipids extracted from BSF larvae can serve as a feedstock for biodiesel production.^5, 6^ These lipids can likely be used as a drop-in biofuel for insertion into existing refinery infrastructure, removing the need to develop expensive new refineries or greatly modify existing ones.^7^ This dual capability of BSF, both as waste managers and biofuel precursors, underscores their versatility in contributing to sustainable and circular practices in multiple industries.

Genetically modifying BSF offers great potential for improvement in all the areas they are currently being utilised and is highly desired. The introduction of transgenic enzymes could enable BSF larvae to effectively metabolise a broader range of organic waste components, degrade pollutants within waste streams, and produce high value biomolecules. Proof of concept for engineering insects for these purposes was demonstrated in our previous studies using transgenic *Drosophila melanogaster*. In one paper we engineered strains to produce a fungal laccase enzyme which successfully degraded industrial pollutants such as BPA *in vivo and in vitro*.^8^ In another paper we generated strains to express organomercurial lyase and mercuric reductase which successfully bioremediated methylmercury into volatile Hg^0^ where it could be easily collected.^9^ When considering the use of BSF as a component of animal feed, they are already being manipulated through breeding programs and simple gene knockouts.^10^ Among the various goals are increasing lipid content, increasing larvae size, and reducing certain components such as chitin that aren’t beneficial in the feed.^11^ Fine tuning the BSF larvae lipid profile, or boosting the overall production of lipids to convert more of the biomass into lipids are also distinct possibilities when considering their application as a biofuel feedstock.

Despite the many exciting opportunities for engineering BSF, the genetic engineering toolkit for BSF is sparse. Basic mutagenesis has been performed with various knockouts^10^, but so far only two papers have been published on transgenic methods for BSF^12, 13^. These papers made use of the *piggyBac* transposon system which has been shown to be an efficient method of generating transgenic lines in *D. melanogaster*^14^ and many other species. Their transgene integration methods involved injections of a donor plasmid and a helper plasmid for expression of the piggyBac transposase, Kou *et al*. outcrossed all G0 injected BSF and screening for a fluorescent marker in the G1 larvae whilst Gunther *et al*. outcrossed only the DsRed positive G0 larvae.

The complete life cycle of the BSF is approximately 36 days, with an additional 4+ days waiting for the G1 larvae to hatch.^15^ Here, we generated transgenic lines by utilising piggyBac mRNA instead and utilised a pre-screening method in the G0 larvae, less than 2 weeks after injection. This allowed us to rapidly determine whether an injection was successful and reduce the significant burden of resources and time that goes into maintaining BSF beyond the pre-screening stage.

## Results

To assay the effectiveness of creating transgenic BSF lines, the *piggyBac* donor plasmid pBXL[pIE1hr5-DsRedT3] shown in **Figure 1(A)** was used. This contains the fluorescent reporter DsRed under the control of the hr5-IE1 promoter. hr5-IE1 is a ubiquitously expressed promoter which is a fusion of the *hr5* enhancer and *immediate early 1* promoter from *Autographa californica nucleopolyhedrovirus* and has been well characterised in other insects.^16^

**Figure 1.**
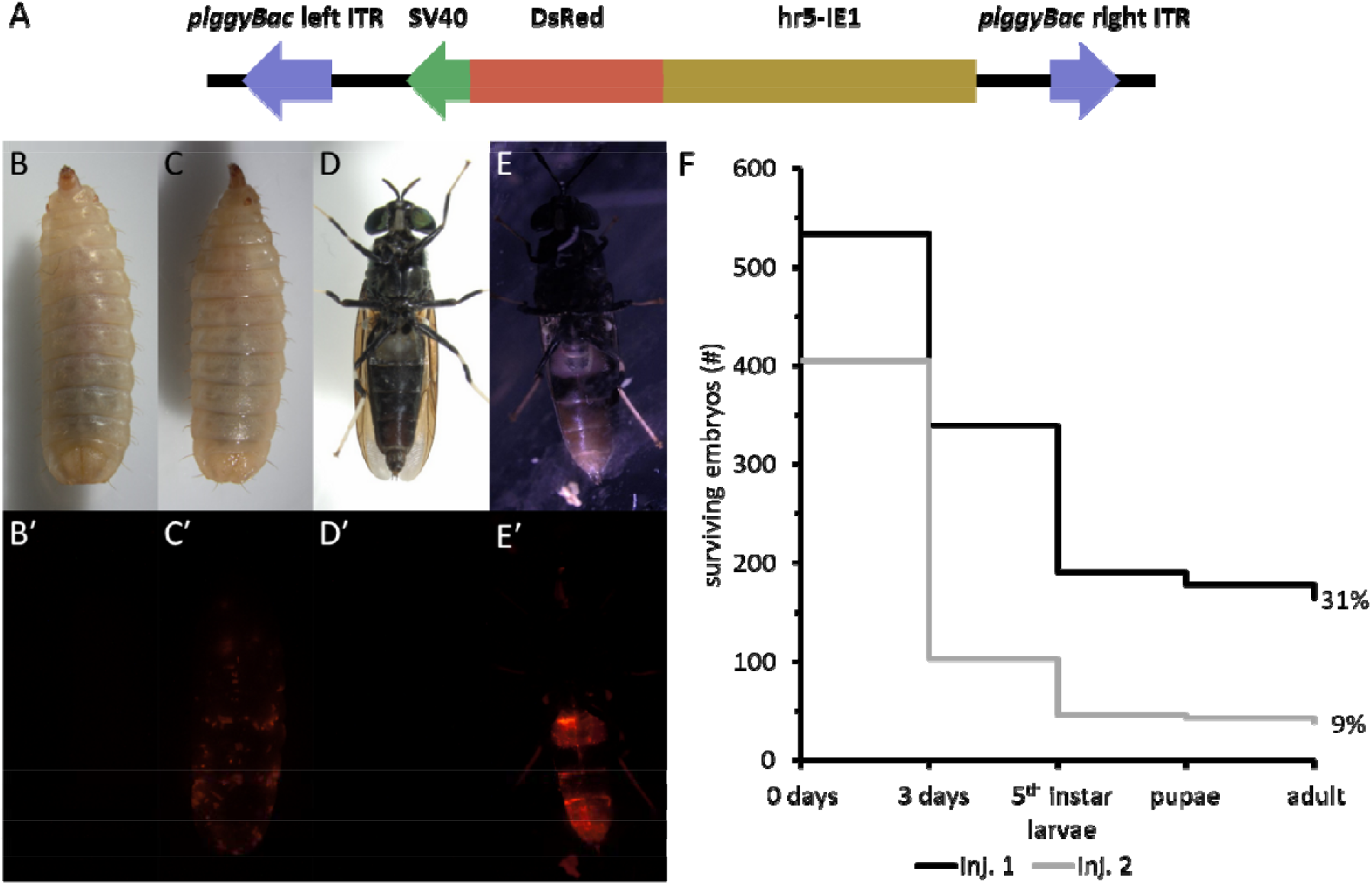
**(A)** Schematic of relevant sections of plasmid pBXL[pIE1hr5-DsRedT3] used to generate transgenic BSF. **(B,C)** Representative bright-field images of the dorsal view of 5^th^ instar BSF **(B)** WT larva and **(C)** G0 mosaic larva. **(D,E)** Representative bright-field images of the ventral view of adult BSF **(D)** WT and **(E)** G0 mosaic. **(B⍰-E⍰)** Matching fluorescent images of **(B-E)** showing DsRed expression. **(F)** Survival of BSF eggs injected with pBXL[pIE1hr5-DsRedT3] for two injection sessions. 0 days shows count of total eggs injected, 3 days shows count of eggs which developed eye spots. Data is from 6 and 8 independent egg clutches for injection session 1 and 2 respectively. Percentage survival is also shown. ITR, inverted terminal repeat; SV40, simian virus 40 polyadenylation signal; hr5-IE1, hr5 enhancer and *immediate early 1* promoter; inj. 1/inj. 2, injection session 1/2; WT, wild type; BSF, black soldier fly.

The BSF embryos were injected less than 4 h after oviposition with a mixture of donor plasmid and piggyBac mRNA. Survival of the injected embryos was assayed at multiple time points for two independent injection sessions as per **Figure 1(F)**. The survival of injected embryos was comparable to observations in other insect species with established microinjection protocols^17, 18^, with 31% and 9% of embryos reaching the adult stage.

To get an early indication of whether transgenic BSF were generated or not, 12-day-old 5^th^ instar larvae were screened for DsRed fluorescence. A range of mosaicism was seen, **Figure 1(C⍰)** shows a representative fluorescent image of a G0 larva with mosaic expression of DsRed in comparison to a wild type (WT) larva shown in **Figure 1(B⍰)**. **Figure 2(A)** shows the count of injected G0 larvae at this time point, from two injection sessions we generated 110 larvae with mosaic DsRed expression, with a further 128 larvae having no DsRed expression.

**Figure 2.**
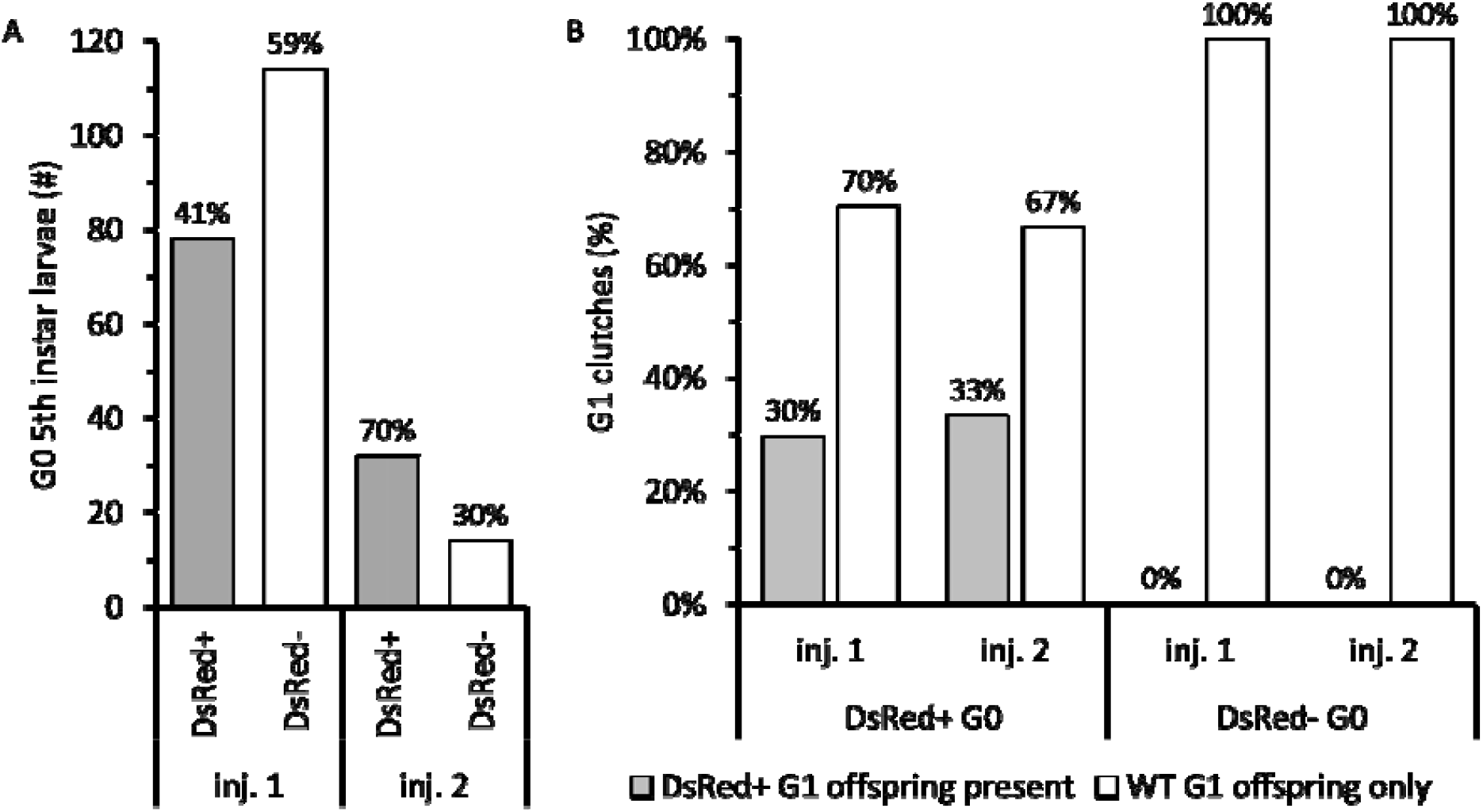
**(A)** Count of G0 5 instar BSF larvae with mosaic DsRed expression vs. no DsRed expression, data is from 14 egg clutches from two injection sessions, totalling 237 larvae. **(B)** Percent of fertilised G1 clutches containing offspring with DsRed expression vs. clutches containing only WT eggs from two injection sessions. Data segmented by DsRed expression of G0 parent. Data is from □58,000 eggs/larvae in 238 egg clutches from injection session 1 and □9,500 eggs/larvae in 36 egg clutches from injection session 2. inj. 1/inj. 2, injection session 1/2; WT, wild type; BSF, black soldier fly.

Mosaic expression of a transgene in G0 animals is no guarantee that the transgene will transmit to the next generation. Likewise, lack of any expression in G0 could be possible if integration occurs in primordial germ cells but not tissues visible when screening G0s for mosaic fluorescence. To investigate this, DsRed-positive and negative G0 pupae were separated into individual vials, sexed after eclosion, and then outcrossed to WT BSF with an approximately 1:2 ratio of G0 mosaics to WT of the opposite sex.

Eggs were collected daily from the outcrossed G0 BSF for 15 days and screened on days 3 and 4 during late egg development or immediately after hatching. The number of clutches containing DsRed-positive offspring vs. the number containing only DsRed-negative offspring for the two injection sessions is shown in **Figure 2(B)**. This data comes from □58,000 eggs/larvae in 238 egg clutches from injection session 1 and □9,500 eggs/larvae in 36 egg clutches from injection session 2. A further 304 egg clutches (□49,000 eggs) across the two injection sessions did not develop and were considered unfertilised, only clutches with >10 fertilised eggs were considered fertilised for screening. DsRed-positive G0’s from injection session 1 produced 27 clutches (29.7%) containing DsRed-positive offspring, similarly DsRed-positive G0’s from injection session 2 produced 9 clutches (33.3%) containing DsRed-positive offspring. No DsRed-negative G0 parents from either injection session produced DsRed-positive offspring indicating that pre-screening G0 parents for mosaic expression is worthwhile. In total, irrespective of DsRed expression in G0’s, 27 of 238 (11.3%) and 9 of 36 (25%) clutches had DsRed-positive offspring from injection sessions 1 and 2 respectively.

The above data for G0 DsRed-positive larvae from injection session 2 was further segmented by the level of mosaicism. Most G0’s had DsRed expression in less than 25% of the whole larvae, 2 females had >25% expression and were crossed to WT separately from the other females. All 3 egg clutches from this cross contained DsRed-positive offspring.

The number of DsRed-positive G1 offspring in any individual clutch that contained DsRed-positive offspring was consistent between injection sessions, 0.7-17.4% for session 1 and 0.6-18.4% for session 2. There were 276 and 120 DsRed-positive G1 offspring in total for injection sessions 1 and 2 respectively.

The G1 screening method above was able to be utilised due to discovering exactly when the hr5-IE1 promoter turns on during development to express DsRed. **Figure 3** provides comparative brightfield and fluorescent images of heterozygous transgenic BSF and WT BSF. **Figure 3(A⍰)** shows a representative image of the late egg stage approximately 3-4 days post-oviposition. It demonstrates the earliest stages of DsRed expression that starts as 1 or 2 patches in the posterior of the developing egg. The larvae that hatched from that same egg is then shown, less than 1 day later in **Figure 3(B⍰)**, by which point the DsRed is being expressed ubiquitously. DsRed expression in later stages of development of heterozygotes is also shown in **Figure 3(C⍰,D⍰)**. Transgene transmission was stable through multiple generations including in backcrosses to make a homozygous line. G4 homozygotes are shown in **Figure 3(I,J)** demonstrating a change in colour under visible light due to the high levels of DsRed expression.

**Figure 3.**
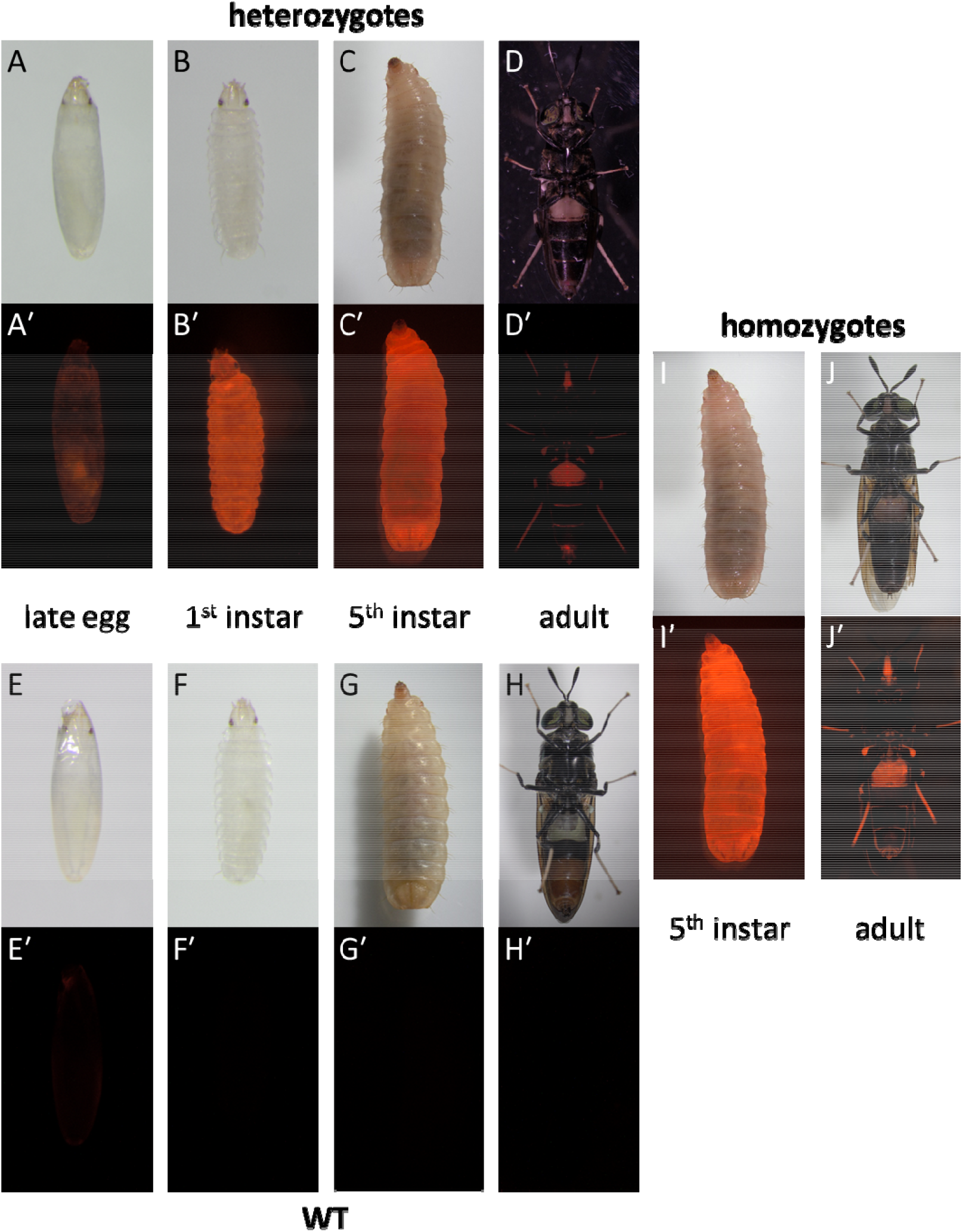
Representative bright-field images of **(A-D)** heterozygous G1 BSF, **(E-H)** WT BSF, and **(I-J)** homozygous G4 BSF at the annotated stage of development. **(A⍰-J⍰)** Matching fluorescent images showing DsRed expression. G1 BSF in **(A)** and **(B)** are the same animal, WT BSF in **(E)** and **(F)** are the same animal. WT, wild type; BSF, black soldier fly.

The location of one transgenic integration was identified by making use of the splinkerette PCR protocol. This method had previously been established to be a quick and economical method for locating piggyBac integrations in *D. melanogaster*.^19^ Digesting the DNA using BstYI was ineffective for BSF as it only yielded 23-37 bp of sequencing in the genome. BglII proved to be more useful and the transgene for one integration was located within a 23 kb intron of the *U-scoloptoxin(01)-Er1a* gene where no deleterious effects were observed.

## Discussion

We have shown that piggyBac mRNA microinjections provide a highly efficient method for generating stable transgenic BSF lines, with good survival after injection and can be screened reliably and quickly in the G0 and G1 generations. Two injection sessions afforded the generation of 396 individual G1 heterozygous transgenic BSF which far exceeds the needed amount to produce multiple stable colonies. Due to positional effects from random integration methods, typically 3-4 independent transgenic lines are used to research the function of the transgene. Based on these numbers, for high throughput applications, where a large number of different transgenic lines are needed, 3-4 different constructs could be injected in a single day and after □ 2 weeks a reliable indicator of success is provided by screening for mosaic expression of a reporter gene.

The previously reported rate of germline transmission was 10 out of 84 (11.9%), this transformation rate was a measure of the percentage of fertile G0 transgenic adults transmitting the transgene to G1 offspring^12^. No data on which individual G0 produced transgenic offspring was recorded in our study due to large group matings and the inability to determine which egg clutches came from which G0 parents. However, if we consider that all fertilised clutches that we collected were only laid from matings between fertile adults, the percentage of those clutches containing transgenic offspring is an indirect measure of the transformation rate. Considering this, our data showed a similar transformation rate of 11.3% for injection session 1 and a substantially higher transformation rate of 25% for injection session 2.

For male G0 transgenics, there is the possibility that some mated with multiple females to produce multiple fertile clutches, and although females will only mate once, they can also produce multiple fertile clutches. This is unlikely to skew the data unless transmitting and non-transmitting G0 adults weren’t producing multiple clutches at the same rate, perhaps due to a fitness cost imposed by the overexpression of DsRed.

Since we have shown that DsRed-negative G0 do not transmit and can be discarded, if we only consider the DsRed-positive G0 BSF, the transformation rate jumps up to 29.7% and 33.3%, showing consistency across the two injection sessions. The 11.9% transmission rate from Kou *et al*. was with the use of a helper plasmid containing the hyperactive piggyBac transposase. This was a vast improvement over the 2.0% transmission rate they achieved when using the non-hyperactive piggyBac transposase helper plasmid. Considering this, use of the hyperactive piggyBac transposase mRNA may see a similarly large increase in transmission rate from what we saw.

The discrepancy of the overall transformation rates in the two injection sessions (11.3% and 25%) vs. the consistency of the DsRed-positive G0 transformation rates (29.7% and 33.3%) can be explained by the lack of transmission from DsRed-negative G0’s. As per **Figure 2(A)**, injection session 1 had no DsRed expression from 59% of the G0’s, giving a subsequently low overall transformation rate. Comparatively, injection session 2 had no DsRed expression from a much lower 30% of G0’s, giving a much higher overall transformation rate.

It is probable that the level of DsRed expression in G0 mosaics correlates with the likelihood of transmission as indicated by the G0’s with >25% expression level transmitting the transgene to offspring in all of their clutches. This was a very small sample size however, but if it does hold up, it will streamline the process further by prioritising the screening of G0 crosses by their level of mosaicism.

To assist with screening, besides G0 mosaicism, our discovery that the hr5-IE1 promoter turns on in the late egg stage of development greatly reduced the complexity and work required for screening G1 eggs and larvae. If this had occurred any later the hatched larvae would need to be put into Gainesville diet where their small size makes it very hard to separate them from the diet for screening. To effectively screen after this point without losing any larvae or without a large amount of work tediously separating small larvae from the diet, you would need to wait a further □2 weeks until they’re 5^th^ instar larvae and big enough to separate from the diet easily. An alternative might have been to keep them out of the feed longer, but that likely would have been detrimental to their health and survival. Additionally, by screening eggs and larvae before being placed into the diet, the transgenic larvae can be separated from WT siblings and not have to compete for survival and could be placed into a smaller container.

Future work has the potential to streamline this process further. A promoter that allows screening earlier than day 3 would mean a wider screening window where transgenic eggs could be separated from WT eggs well before hatching.

The prior research had cast doubt on the effectiveness of mRNA injections, attributing failures to RNAse activity. As has been shown however, our data reveals this is not the case. This now paves the way for exploring alternative mRNA constructs and their applications in genome editing BSF through microinjection.

Locating the piggyBac integration site was achieved by adapting the previously published splinkerette PCR protocol^19^. This has proven to be a rapid and simple procedure which only requires Sanger sequencing so can be done economically. This will also reduce the complexity of screening for and generating homozygous vs heterozygous transgenic lines and has the potential to identify safe harbour sites for more targeted transgenic integration methods.

## Conclusions

In summary, our study demonstrates the efficacy of mRNA injections as a successful method for generating transgenic BSF. When combined with our pre-screening method, this allows for the rapid and efficient transmission of transgenes to successive generations. Looking ahead, our research will focus on expanding the toolkit of transgenic techniques and testing a diverse suite of endogenous BSF promoters to enable precise control over transgene expression in different tissues and life stages. These efforts will allow fine tuning of BSF bioproduction to better suit the practical applications and to thrive on a wider variety of waste, further fostering advancements in both basic and translational research.

## Methods

The aim of this study was to assess the effectiveness of generating stable transgenic BSF via the use of piggyBac mRNA microinjections, and to determine efficient ways of screening for success.

Standard manufacturer’s protocols were used for all commercial kits and reagents unless otherwise stated.

## Plasmids

PiggyBac donor plasmid pBXL[pIE1hr5-DsRedT3] was a gift from Alfred Handler in the Center for Medical, Agricultural and Veterinary Entomology at the USDA Agricultural Research Service (USDA-ARS-CMAVE).

Plasmids were transformed into *E. coli* Stbl3 cells, miniprepped (NEB Monarch Plasmid Miniprep Kit), sequence verified, and midiprepped (Macherey-Nagel NucleoBond Xtra Midi kit) for microinjection.

## mRNA generation

dsDNA for the transcription of mRNA was amplified (NEB Q5 High-Fidelity DNA Polymerase) from phsp-pBac^20^ using primers GAAACTAATACGACTCACTATAGGGAGAGCCGCCACatgggtagttctttagacgatg and TCAGAAACAACTTTGGCACATATCA in a total volume of 125 μL with a 62 °C annealing temperature and 55 s extension time. The PCR product was purified using paramagnetic beads (Beckman Coulter RNAClean XP Reagent).

The IVT reaction was performed (NEB HiScribe T7 ARCA mRNA Kit (with tailing)) for 16 h. An additional 5 μL 2X NTPs (NEB 2X ARCA/NTP Mix) was added to the tailing reaction which was run for 1 h. Final product was purified using paramagnetic beads (Beckman Coulter RNAClean XP Reagent). See Additional File 1 for detailed protocol.

## BSF husbandry

The BSF Dornoch strain was used for all experiments in this manuscript. This strain was sourced from wild BSF in the Brisbane (Australia) urban area.

BSF were reared in a contained environment room (CER) at 26 °C and 70% relative humidity. CER was lit under a 13 h light and 10 h dark cycle with 30 min transition period between each. Sunrise was set to 5 AM to facilitate egg laying during working hours of the day. Additional LED lights were added directly adjacent to or above the cages to enhance fertility.

Adult flies were kept in cube-shaped 32.5 cm or 47.5 cm cages (BugDorm 4M Series Insect Cages) and provided with water bottles containing wicks for drinking. 120 mL plastic cups of D□30 %V/V honey (Cloverdale Pure Honey) in water were also placed in the cages with paper towel scrunched up into the honey water to prevent direct access to it.

Adult males and females were placed in the same cages for group mating and a cave was set up for egg-laying. It consisted of a 120 mL plastic cup containing 12 g poultry pellets mixed with 100 mL water. The plastic cup was placed in a 500 mL plastic food container which already had a 50:50 mixture of apple cider vinegar (Cornwell’s Apple Cider Vinegar) and apple juice (Woolworths Apple Juice) along with several dead adult BSF. The plastic food container was sealed with micropore tape (3M Micropore Surgical Tape, 1530-3). Two terracotta tiles with □1.5 mm in-cut grooves were stacked on top of each other and placed on top of the plastic food container. A black pot plant pot with a window cut in the side was placed over the entire thing, upside down. Eggs were collected either the same day for injections and screening, or 2-3 days later for colony maintenance.

Freshly hatched eggs were put into Gainesville diet immediately or after 3-4 days if screening was required. Gainesville diet was topped up or replaced with fresh diet if the top layer dried out or grew mould.

## Generation of transgenic lines

Microinjection needles were pulled from quartz capillaries (Sutter Instrument quartz capillaries, QF100-70-10) using a needle puller (Sutter Instrument Model P-2000) with the following parameters: Heat 735, Fil 4, Vel 42, Del 126, and Pul 160. Pulled capillaries were then ground on a microgrinder (Narishige EG-45 Microgrinder), creating as small an opening as possible. 5 μL of microinjection mixture was prepared on ice containing final concentrations of 300 ng/μL donor plasmid and 300 ng/μL piggyBac mRNA, with 0.5 μL green dye (Queen Classic Green Food Colour) filtered at 0.2 μm (Sarstedt 0.2 μm syringe filter, 83.1826.001). All microinjection reagents were prepared in RNAse free conditions.

Egg collection tiles were placed in adult BSF cages mid-morning and eggs were collected from □30 min and onwards after placing the tile. Eggs were brushed onto moist filter paper with a fine brush and aligned parallel to one another with the posterior end against the edge of a glass coverslip. Coverslip was removed and a second coverslip was half-covered in tape (Double-sided Sellotape) before being gently lowered onto the aligned eggs using tweezers so they attached with posterior ends facing outward. Eggs were covered with halocarbon oil (Sigma-Aldrich Halocarbon oil 700). Under 80x magnification on a stereo microscope, eggs were injected (Eppendorf FemtoJet 4i/Injectman 4) with the smallest amount of visible microinjection mixture using a combination of constant compensation pressure and single injection pressures in the range of 5-15 psi depending on effectiveness. Injections were performed with the needle entering into the posterior of the eggs at a horizontal angle.

After injection of all eggs on a single coverslip, halocarbon oil was washed off with de-ionised water under gentle pressure from a squeeze bottle before being placed in the CER for 3 days. After screening for eye spots at day 3, eggs were transferred to Gainesville diet until □day 12 (5^th^ instar larvae) before being screened for fluorescence. Larvae were then returned to Gainesville diet for □2 weeks next to a secondary container of wood shavings (Peckish Pet Bedding Classic) before separating all pupated larvae into individual vials (Genesee Scientific Narrow Drosophila Vials, 32-109RL) and sealing with flugs (Genesee Scientific Flugs, 076-49-102). After eclosion, adult flies were sexed and separated into 2 cages of males and females with mating caves set up as described above. □2x WT adult virgin flies of the opposite sex were added to the cages. As more G0 flies eclosed in the following weeks, they were added to the same cages and additional WT flies were added at the same time.

Each day after the mating cages were set up, the tiles were replaced, and individual egg clutches were transferred to petri dishes (Greiner 60×15 mm petri dishes, 628161). 2- and 3-day old eggs were screened for eye spots, if present, petri dishes were covered in parafilm with several holes punctured in it, then sealed with micropore tape (3M Micropore Surgical Tape, 1530-3). Clutches with eye spots were then screened again 1-2 days later when they had hatched or just before hatching for fluorescence. If eggs had no eye spots by day 4, they were considered unfertilised eggs.

## Locating transgenes

5^th^ instar larvae were placed in -30 °C for 2 days then anterior segments 1 to 3 were excised with a scalpel to be used for DNA extraction. DNA was extracted (Roche High Pure PCR Template Preparation Kit) following the kit instructions for mouse tail DNA.

The splinkerette PCR protocol was used to locate the transgenes.^19^ 750 ng of DNA was digested with BglII (NEB BglII), purified using Qiagen PCR Purification kit, and a modified 3’SPLNK-PB-SEQ primer was used due to a SNP in our piggyBac inverted terminal repeat (ACaCATGATTATCTTTAAC).

## Supporting information

Additional File 1 - Protocol

## List of abbreviations

BPA: bisphenol A
BSF: black soldier fly
CER: contained environment room
*D. melanogaster*: *Drosophila melanogaster*
DNA: deoxyribonucleic acid
dsDNA: double-stranded deoxyribonucleic acid
DsRed: *Discosoma sp*. red fluorescent protein
*E. coli*: *Escherichia coli*
G0: generation 0
G1: generation 1
G4: generation 4
hr5-IE1: *hr5* enhancer and *immediate early 1* promoter
inj.: injection
mRNA: messenger ribonucleic acid
NTPs: nucleoside triphosphates
PCR: polymerase chain reaction
RNase: ribonuclease
SV40: simian virus 40 polyadenylation signal
SNP: single nucleotide polymorphism
WT: wild type.

## Declarations

### Ethics approval and consent to participate

Not applicable.

### Consent for publication

Not applicable.

### Availability of data and materials

All data generated or analysed during this study are included in this published article.

### Competing interests

MM and CP are co-founders of EntoZyme PTY LTD.

### Funding

Funding for this study was provided by Fit Milestones Pty Ltd.

### Authors’ contributions

MM conceived of the study and provided guidance and supervision to other authors. RAH and CP developed the microinjection protocols. SK prepared the plasmids. CP and KT generated the mRNA. CP, CR, JMM, KT, and SK performed the BSF husbandry. CP and KT generated the transgenic lines. CP screened the G0 mosaic BSF. CP, CR, JMM, KT, MM, and SK screened the G1 BSF. CP located the transgenes. CP wrote the manuscript and created the figures. All authors read and contributed to the final manuscript.

## Acknowledgements

The authors are grateful to Dr. Cate Paull at CSIRO for her kind gift of *H. illucens* Dornoch strain and the initial guidance on BSF rearing protocols.

## Additional files

Additional File 1.pdf

Title: PiggyBac mRNA Production

Description: A detailed protocol for the generation and purification of the piggyBac mRNA used in this study.

## Notes

### Competing Interest Statement

CP and MM have commercial interests in EntoZyme PTY LTD

