## Additional File 1 - Protocol for "Rapid generation and screening of transgenic black soldier fly (*Hermetia illucens*)"

### PiggyBac mRNA Generation

#### Reagents required

- Agarose
- Beckman Coulter RNAClean XP, A63987
- 100% ethanol (RNase-free)
- MQ $H_2O$ (RNase-free)
- NEB Deoxynucleotide (dNTP) Solution Mix, N0447
- NEB DNase I (RNase-free), M0303
- NEB HiScribe® T7 ARCA mRNA Kit (with tailing), E2060S
- NEB Q5® High-Fidelity DNA Polymerase, M0491
- NEB RNA Loading Dye, (2X), B0363S
- pBac forward primer
    - `5'-GAAACTAATACGACTCACTATAGGGAGAGCCGCCACatgggtagttctttagacgatg-3'`
- pBac reverse primer
    - `5'- TCAGAAACAACTTTGGCACATATCA-3'`
- phsp-pBac plasmid
- ThermoFisher RiboRuler High Range RNA Ladder, SM1821

#### Equipment required

- Bio-Rad 16-Tube SureBeads™ Magnetic Rack, # 1614916
- BSL cabinet/sterile hood
- Gel electrophoresis equipment
- NanoDrop machine
- Thermocycler

#### Notes

After completing the PCR, all work should be done under RNAse-free conditions, in a BSL cabinet/hood that can be decontaminated with UV light beforehand. Gel electrophoresis tank, gel mould and comb should all be sterilised with NaOH pellets or some other method before running RNA samples.

#### Protocol

1. Mix the following 9.5x master mix reagents and aliquot into 8 PCR tubes:

| **Reagent** | **1x (µL)** | **8.5x MM (µL)** |
|---|---|---|
| NEB 5X Q5 Reaction Buffer (5x) | 5 | 42.5 |
| pBac forward primer (10 µM) | 1.25 | 10.625 |
| pBac reverse primer (10 µM) | 1.25 | 10.625 |
| NEB Deoxynucleotide (dNTP) Solution Mix (10 mM) | 0.5 | 4.25 |
| NEB Q5 High-Fidelity DNA Polymerase (2 U/µL) | 0.25 | 2.125 |
| MQ $H_2O$ | 15.75 | 133.875 |
| **Total** | **24** | |

2. Add 1 µL of phsp-pBac plasmid (1 ng/µL) to 7 tubes and 1 µL MQ $H_2O$ to the final tube.

3. Place the tubes in a thermocycler with the following parameters:

| | Temperature | Time |
|---|---|---|
| 1 | 98 °C | 30 s |
| 2 | 98 °C | 10 s |
| 3 | 62 °C | 30 s |
| 4 | 72 °C | 55 s |
| 5 | Go to step 2 | 34 times |
| 6 | 72 °C | 2 min |
| 7 | 4 °C | ∞ |

4. Make a 1% agarose gel and run 5 µL of each PCR product for 50 min @ 100 V. A Band should be present at 1821 bp.
5. Combine all plasmid PCR reactions with the correct band into a 1.5 mL tube to make a 140 µL solution.
6. Gently shake the RNAClean XP bottle, resuspending the settled magnetic particles.
7. Add a 252 µL RNAClean XP and mix up and down with pipette 15 times. Solution should be homogenous after mixing.
8. Incubate 5 min @ RT.
9. Place tube onto magnetic rack until solution is clear (~5 min).
10. Make 70% ethanol using 1400 µL nuclease-free ethanol and 600 µL nuclease-free MQ $H_2O$.
11. Gently remove and discard cleared solution with a pipette, avoiding contact with magnetic beads.
12. Add 500 µL of the 70% ethanol, the ethanol must completely cover the bead mass on the side of the tube.
13. Incubate 30 s @ RT.
14. Gently remove the ethanol with a pipette and repeat the previous 2 steps 2 more times. When removing the ethanol for the final time, do it initially with a P1000, then P100, then P2 to remove as much as possible.
15. Incubate @ RT in hood with lid open until sample completely dries (~10 min).
16. Remove tube from magnetic rack.
17. Resuspend bead pellet in 30 µL MQ $H_2O$ by pipetting up and down several times after washing all the magnetic beads into solution.
18. Place on magnetic rack until solution is clear (~5 min).
19. Gently transfer cleared solution with pipette to a clean 1.5 mL tube, avoiding contact with magnetic beads.
20. Check concentration of this purified piggyBac PCR on Nanodrop is above 250 ng/µL and free of contamination.

21. Thoroughly mix the following reagents in 2x PCR tubes:

| Reagent | Amount |
|---|---|
| Nuclease-free MQ $H_2O$ | up to 20 µL |
| NEB 2X ARCA/NTP Mix (2x) | 10 µL |
| Purified PiggyBac PCR product | 1000 ng |
| NEB T7 RNA Polymerase Mix | 2 µL |
| **Total** | **20 µL** |

22. Incubate tube 16 h @ 37 °C in thermocycler.
23. Transfer 2 µL to PCR tube on ice for testing later.
24. Add 4 µL NEB DNase I (RNase-free) to tube and mix well.
25. Incubate 15 min @ 37 °C.
26. Transfer 2 µL to PCR tube on ice for testing later.
27. Mix the following reagents in a PCR tube:

| Reagent | Amount (µL) |
|---|---|
| Nuclease-free $H_2O$ | 60 |
| DNase I-treated IVT product | 20 |
| NEB 10x Poly(A) Polymerase Reaction Buffer (10x) | 10 |
| NEB 2X ARCA/NTP Mix (2x) | 5 |
| NEB *E. coli* Poly (A) polymerase (5 U/µL) | 5 |
| **Total** | **100** |

28. Incubate 30 min @ 37 °C.
29. Transfer to a clean 1.5 mL tube.
30. Transfer 2 µL to PCR tube on ice for testing later.
31. Gently shake the RNAClean XP bottle, resuspending settled magnetic particles.
32. Add 176.4 µL RNAClean XP and mix up and down with pipette 15 times. Solution should be homogenous after mixing.
33. Incubate 5 min @ RT.
34. Place tube onto magnetic rack until solution is clear (~5 min).
35. Make 70% ethanol using 1400 µL nuclease-free ethanol and 600 µL nuclease-free MQ $H_2O$.
36. Gently remove and discard cleared solution with a pipette, avoiding contact with magnetic beads.
37. Add ~300 µL of the 70% ethanol, the ethanol must completely cover the bead mass on the side of the tube.
38. Incubate 30 s @ RT.
39. Gently remove the ethanol with a pipette and repeat the previous 2 steps 2 more times. When removing the ethanol for the final time, do it initially with a P1000, then P100, then P2 to remove as much as possible.

40. Incubate @ RT in hood with lid open until sample completely dries (~10 min).
41. Remove tube from magnetic rack.
42. Resuspend bead pellet in 30 µL nuclease-free MQ $H_2O$ by pipetting up and down several times after washing all of the magnetic beads into solution.
43. Place on magnetic rack until solution is clear (~5 min). You may need to take out central strip of magnets from the rack, lay it down and then lay tube on top of it and positioning the liquid above the magnet, then after the 5 minutes slow turn the magnet strip and tube so the liquid rolls down to the bottom of the tube without any beads.
44. Gently transfer cleared solution with pipette to a clean 1.5 mL tube, avoiding transfer of magnetic beads.
45. Transfer 2 µL to PCR tube on ice for testing.
46. Check concentration on Nanodrop.
47. Add 2 µL NEB RNA Loading Dye, (2X) to each of the 2 µL RNA aliquots set aside.
48. Add 4 µL ThermoFisher RiboRuler High Range RNA Ladder and 4 µL NEB RNA Loading Dye, (2X) to a clean PCR tube on ice.
49. Place all those mixtures in a thermocycler at 70 °C for 10 min.
50. Place tubes back on ice.
51. Make an RNase-free, 1.5% agarose gel and run RNA aliquots 80 min @ 100 V with Ladder Riboruler High Range. Pre-tailed band should be present at 1785 bp and tailed band should be higher.
52. Store at -80 °C.
